# Drug promiscuity: problems and promises

**DOI:** 10.1101/2023.12.29.573633

**Authors:** Antonius M. VanDongen

**Affiliations:** Department of Pharmacology and Cancer Biology, Duke University, Durham, North Carolina, USA

## Abstract

Therapeutic drugs exert their effect by binding to a specific target protein (receptor) and altering its activity. However, all drugs have undesirable side effects, which likely result from the drug binding to other protein targets. Drug promiscuity refers to the ability of small molecule compounds to bind with high affinity to multiple proteins. This drug property is responsible for the problems caused by their side effects. But drug promiscuity also presents a promise: if a drug targets numerous proteins, each interaction may contribute to its therapeutic efficacy. The ability to target multiple genes is especially promising for medicines that treat systemic diseases, including neurological disorders and cancers, which result from gene network problems. Here, I will review both aspects of drug promiscuity and illustrate a potential strategy to move this forward using a computer modeling approach.

## Introduction

Therapeutic drugs are designed to achieve their medicinal effects primarily through specific binding to target proteins. This interaction is critical in modulating protein activities, either inhibiting or altering their functions, and forms the basis of drug efficacy in treating a myriad of diseases. However, a common and often problematic aspect of these drugs is their unintended side effects. These adverse effects are frequently attributed to the drug’s interaction with proteins other than the intended target.

Drug promiscuity, defined as the ability of a single drug molecule to interact with multiple protein targets, presents a paradox in pharmacology. It is a source of therapeutic potential and unwanted side effects, making its study challenging and essential. While it often leads to undesirable side effects, this property also holds potential for therapeutic innovation. Many drugs, being small molecule compounds, have the capacity to bind with high affinity to several different proteins. While this can cause unwanted reactions, it also means that a single drug can have a multi-targeted approach, potentially increasing its therapeutic value.

Unfortunately, the specific targets of many approved drugs remain unidentified, and the molecular mechanisms underlying their side effects are even less understood. This knowledge gap represents a significant challenge in drug development and safety. However, drug promiscuity also presents a unique opportunity. Suppose we can unravel the complex gene networks involved in systemic diseases, such as neurological disorders and cancers. In that case, we may be able to identify drugs that selectively target a subset of proteins within these networks. This approach could enhance drug efficacy while minimizing side effects.

In this context, I aim to review both the challenges and opportunities presented by drug promiscuity. I will explore how an in-silico approach – using computer simulations and models – could be a valuable strategy for understanding and harnessing this phenomenon. We can devise more effective and safer therapeutic strategies by computationally analyzing the interactions between drugs and protein networks. This in-silico methodology represents a promising frontier in drug discovery and development, offering a path to understand better and exploit the complex nature of drug-protein interactions.

## Drug-Receptor Interactions

Drugs exert their medicinal effect by binding to a ‘receptor’ and modulating its function. These receptors are typically cellular proteins that play a critical role in a disease process. Examples include neurotransmitter receptors, ion channels, or enzymes. Drugs physically interact with a receptor by closely associating with a binding site, a part of the receptor protein that is compatible with both the 3-dimensional structure of the drug and its chemical properties. This interaction affects the protein’s function in the cell: occupancy of the binding site by the drug may prevent an endogenous compound from binding, or it can induce (or prevent) a conformational change in the protein. While the receptor is the intended therapeutic target for the drug, it may also bind to other proteins with functions unrelated to the disease process. This kind of promiscuity may cause unwanted side effects. In this manuscript, we will refer to the ability of drugs to bind to multiple proteins as ‘promiscuity.’ It is the opposite of specificity.

## Affinity and Efficacy

Two essential properties characterize drug-receptor interactions: affinity and efficacy. Affinity specifies how ‘tight’ a drug binds to its receptor. Efficacy refers to the ability of a drug to ‘activate’ its receptor. High affinity is a desirable property of a drug, allowing it to have therapeutic effects at a relatively low concentration. Such a strong interaction then helps to minimize unwanted side effects. Drug binding is a dynamic, reversible process in which the drug associates with the receptor, occupies its binding site for some time, and disassociates again. The longer the drug is associated, the higher its affinity. Drugs don’t have to have efficacy to be effective. They can occupy their binding site without activating the receptor, thereby preventing a natural compound (e.g., a neurotransmitter) or a protein segment (e.g., enzyme-substrate) from binding. This type of interference is the basis for competitive antagonism. Alternatively, a drug could bind at a site away from the natural ligand binding site and allosterically modulate receptor function by inducing or preventing a conformational change.

## Drug promiscuity

I have performed a relatively small-scale study of drug promiscuity using a computational approach by inverting the process of *in silico* drug discovery, in which an extensive library of small molecules is “docked” into a binding site of a protein of interest. Docking involves finding the optimal pose (three-dimensional fit) of each drug in the binding site and estimating its affinity using a force field model that parameterizes all the chemical and physical interactions between the drug and the binding site. The top-ranking drugs with the highest estimated affinity can be further characterized by appropriate assays to evaluate their efficacy.

Here, we would like to do the opposite: dock each drug of interest into a library of protein binding sites. Then, rank the proteins by the estimated affinities of the drug for each binding site. Promiscuous drugs will rank high for many proteins, while an ideal specific drug will only bind with high affinity to a single target. Commercial docking software is not designed to work this way. But that is not a problem: once we have identified the library of proteins and a list of drugs, we dock all drugs into each target binding site, using the standard approach. When we have an estimated affinity and relative rank order for each drug-target combination, we reorganize the data to get the statistics we seek.

## Methods

As of the current analysis, the US Food and Drug Administration (FDA) has approved 2,761 small-molecule drugs. This figure contrasts the estimated 25,000 proteins the human genome can express, highlighting a vast landscape of potential drug-protein interactions. To do this analysis exhaustively, one must perform 70 million dockings. Before embarking on such a significant undertaking, I performed a much smaller-scale analysis to discover what could be learned from such an approach. The protein library was limited to proteins co-crystallized with an FDA-approved drug or a natural hormone or neurotransmitter. This has two advantages: the structures already have a known binding site, and the protein targets (receptors) most likely are therapeutically relevant. I identified 622 receptors in the protein structure database at www.rcsb.org and downloaded their structure data files containing atomic coordinates. **Table 1** shows 20 entries, listing the drug name, PDB identifier, protein/gene name, and therapeutic class/use:

**Table 1.** Examples of PDB entries selected for this study.

| Drug | PDB | Gene/Protein | Therapeutic class/use |
| --- | --- | --- | --- |
| Acetazolamide | 5jn8 | Carbonic anhydrase | Diuretic |
| Alclofenac | 1ht8 | Cox-1 | Non-steroidal anti-inflammatory (NSAID) |
| Acyclovir | 1pwy | Purine Nucleoside Phosphorylase | Anti viral |
| Alendronate | 2f92 | Farnesyl PyroPhosphate Synthase | Treats osteoporosis |
| Alprazolam | 3u5j | BRD4 bromodomain | Psychotropic, anxiolytic |
| amiodarone | 5h4d | Sirtuin 3 (SIRT3) | Antiarrhythmic |
| Amlexanox | 4wbo | GPCR kinase 1 | Treats canker sores (discontinued) |
| Amodiaquine | 2aou | Histamine methyltransferase | Antimalarial |
| Aspirin | 1f9g | Hyaluronate lyase | Salicylate, painkiller |
| Aspirin | 1oxr | Phospholipase A2 | Salicylate, painkiller |
| Aspirin | 2x08 | Cytochrome c peroxidase | Salicylate, painkiller |
| Aspirin | 3tgy | Lactoperoxidase | Salicylate, painkiller |
| Aspirin | 3vln | Glutathione transferase | Salicylate, painkiller |
| Atenolol | 2nuv | Lactoferrin | Beta blocker |
| Atorvastatin | 1hwk | HMG-CoA reductase | Statin |
| Atovaquone | 1uum | Dihydroorotate dehydrogenase | Antiprotozoal agent |
| Atropin | 3osh | Phospholipase A2 | Muscarinic antagonist |
| Azathioprine | 4acs | Glutathione transferase | Immunosuppressant |
| Aztreonam | 3pbs | Penicillin-binding protein 3 | Monobactam antibiotic |

Some drugs, like aspirin in **Table 1**, have been co-crystalized with several distinct proteins, illustrating drug promiscuity. Some proteins, like phospholipase A2 in Table 1, have been co-crystalized with multiple drugs, illustrative of protein promiscuity. This aspect will not be developed in depth in this paper. A total of 1531 small molecules approved as therapeutic drugs by the FDA were downloaded from the ZINC database (www.zinc20.org) as 3D structures in SDF format (Irwin 2012, Irwin 2020). The 1531 FDA-approved drugs were docked in each of the 622 protein-drug complex structures, and their affinities were estimated. Docked drugs were ranked by their affinity for each target, and their rank orders were stored together with the protein target’s PDB identifier and estimated affinity. The eHiTS software was used for docking and affinity estimation (Zsoldos 2006, Zsoldos 2007). A total of 780,000 docks were performed for this small-scale analysis. Because of the large number of docks required, the High Throughput Screening option was used in eHiTS.

## Results

The traditional approach of docking a library of small molecules (FDA drugs) into a target, which was employed here, also generates potentially interesting data unrelated to drug promiscuity. Top-ranking drugs could be studied to see whether their high predicted affinity for a target indicates that they are suitable for drug repurposing. **Figure 1** shows an example. One of the targets analyzed here is carbonic anhydrase II ([1a42.pdb](https://www.rcsb.org/structure/1a42)), which was crystallized in a complex with brinzolamide, a drug used to treat glaucoma. The top-ranking drug from the docking screen was celecoxib (Celebrex), a COX-2 inhibitor and nonsteroidal anti-inflammatory drug (NSAID). Its eHiTS score was very high (-12.4), corresponding to an estimated affinity of 1 pM. **Figure 1** shows the optimal poses for the two drugs.

**Figure 1.**
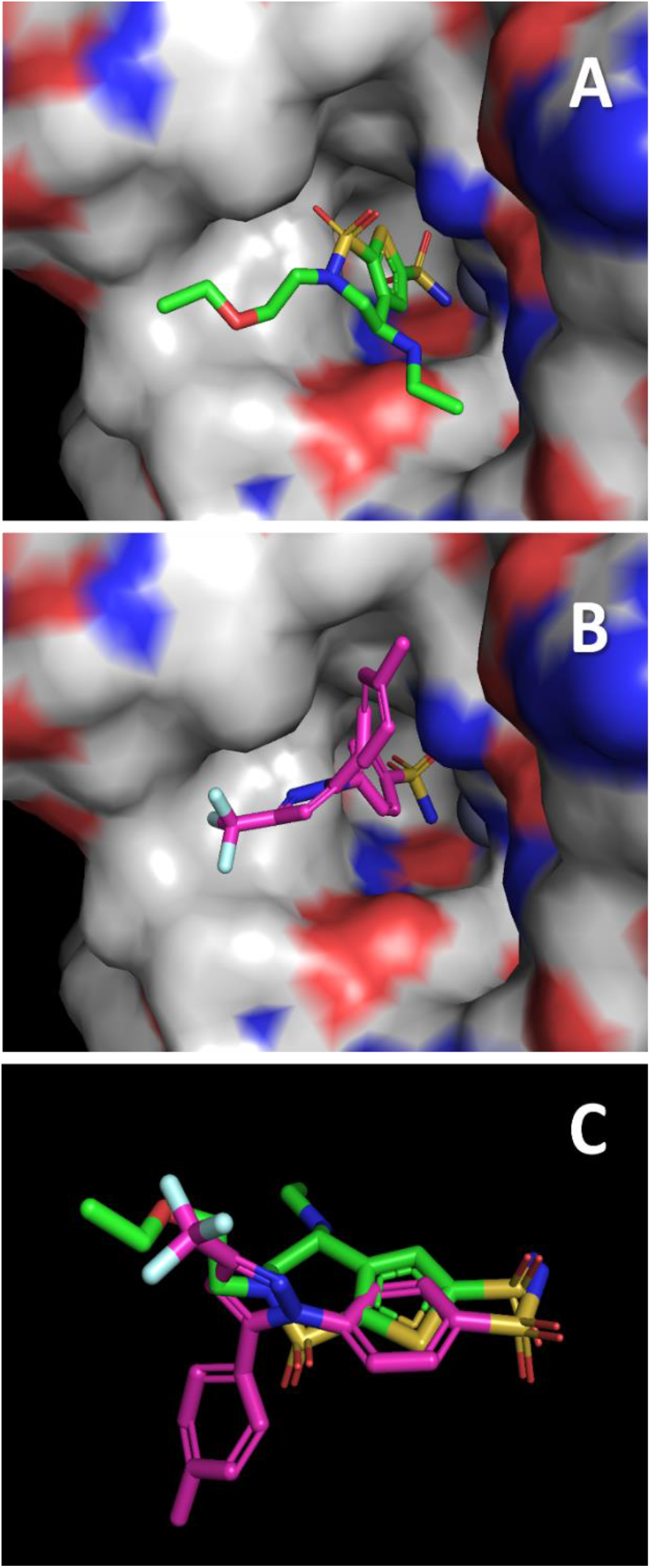
Celecoxib vs Brinzolamide. The binding site for brinzolamide in carbonic anhydrase II is shown in an electrostatic surface representation. **A**. Brinzolamide (sticks) as it was seen in the X-ray structure (1a42.pdb). **B**. The optimal pose for celecoxib in the same binding site, as identified by eHiTS docking. **C**. Comparison of the poses for Brinzolamide (purple) and Celecoxib (green).

**Figure 1.**
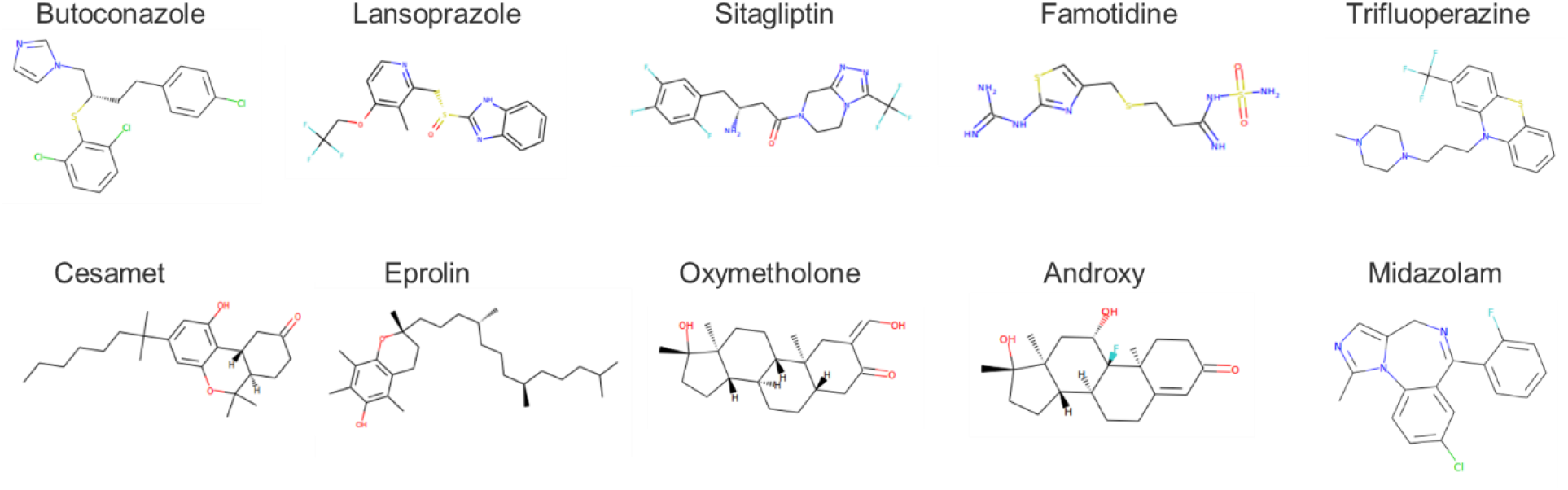
Structures of the ten most promiscuous drugs.

Interestingly, celecoxib has already been shown to be a potent inhibitor of carbonic anhydrase II (Alper 2006). However, it may not be an excellent drug to repurpose for glaucoma treatment because several of its side effects involve problems in the eyes. Side effects are important considerations that often limit the successful repurposing of drugs.

There are probably additional novel drug-receptor matches in this data set, some of which may be useful for drug repurposing. Still, a complete analysis of this topic is outside the scope of this paper. A comprehensive version of the analysis performed here, including all known protein structures and all FDA-approved drugs, could yield a wealth of new receptor-drug interactions.

Now that the rank orders are available for all drug-receptor combinations that were docked, it is possible to organize the data focusing on individual drugs and their rank orders.

For each drug, the rank orders for the 622 receptors were sorted by their estimated affinity, from high to low (**Table 2**). To identify promiscuous drugs, we need to come up with a criterion. I identified the top 1% ranks: because 1531 drugs were competing for each binding site, the top 1% would rank below 15. In **Table 2**, they are highlighted in yellow. Aspirin, baclofen, carteolol, and doxylamine have 1, 2, 2, and 11 better than 1% rankings, respectively. To identify the most promiscuous drugs, I counted the number of ranks below 15 and sorted by counts (**Table 3**).

**Table 2.** Sorted rank orders. Each column lists how the drug ranked relative to all other drugs for a given receptor, sorted from high to low. So, the highest aspirin ranked in any of the 622 receptors was 6, and its next best rank was 22. Rank scores below 15 (top 1%) are highlighted in yellow. Doxylamine scored 11 times below 15.

| drug name | Aspirin | Baclofen | Bethanechol | Carbinoxamine | Carteolol | Cyclopentolate | Diethylpropion | Doxylamine | Esmolol | Ethotoine |
| --- | --- | --- | --- | --- | --- | --- | --- | --- | --- | --- |
| 1 | 6 | 6 | 143 | 73 | 7 | 42 | 64 | 2 | 38 | 15 |
| 2 | 22 | 13 | 181 | 146 | 12 | 74 | 91 | 2 | 42 | 45 |
| 3 | 23 | 13 | 185 | 155 | 21 | 78 | 141 | 2 | 47 | 47 |
| 4 | 26 | 18 | 223 | 159 | 21 | 92 | 163 | 3 | 53 | 61 |
| 5 | 29 | 19 | 228 | 166 | 22 | 96 | 165 | 7 | 57 | 66 |
| 6 | 30 | 21 | 263 | 167 | 24 | 104 | 183 | 9 | 64 | 70 |
| 7 | 35 | 22 | 264 | 177 | 30 | 112 | 192 | 9 | 68 | 74 |
| 8 | 36 | 26 | 267 | 197 | 30 | 118 | 203 | 12 | 73 | 74 |
| 9 | 43 | 27 | 268 | 219 | 35 | 123 | 217 | 12 | 75 | 82 |
| 10 | 43 | 30 | 269 | 221 | 35 | 124 | 243 | 13 | 80 | 85 |
| 11 | 47 | 34 | 283 | 223 | 43 | 130 | 245 | 13 | 84 | 87 |
| 12 | 48 | 36 | 287 | 229 | 46 | 135 | 251 | 15 | 89 | 95 |
| 13 | 51 | 37 | 289 | 237 | 48 | 138 | 264 | 18 | 98 | 100 |
| 14 | 67 | 39 | 303 | 237 | 49 | 139 | 267 | 19 | 102 | 102 |
| 15 | 70 | 41 | 318 | 238 | 50 | 144 | 276 | 19 | 105 | 105 |
| 16 | 72 | 45 | 321 | 255 | 52 | 147 | 281 | 22 | 105 | 110 |

**Table 3.**
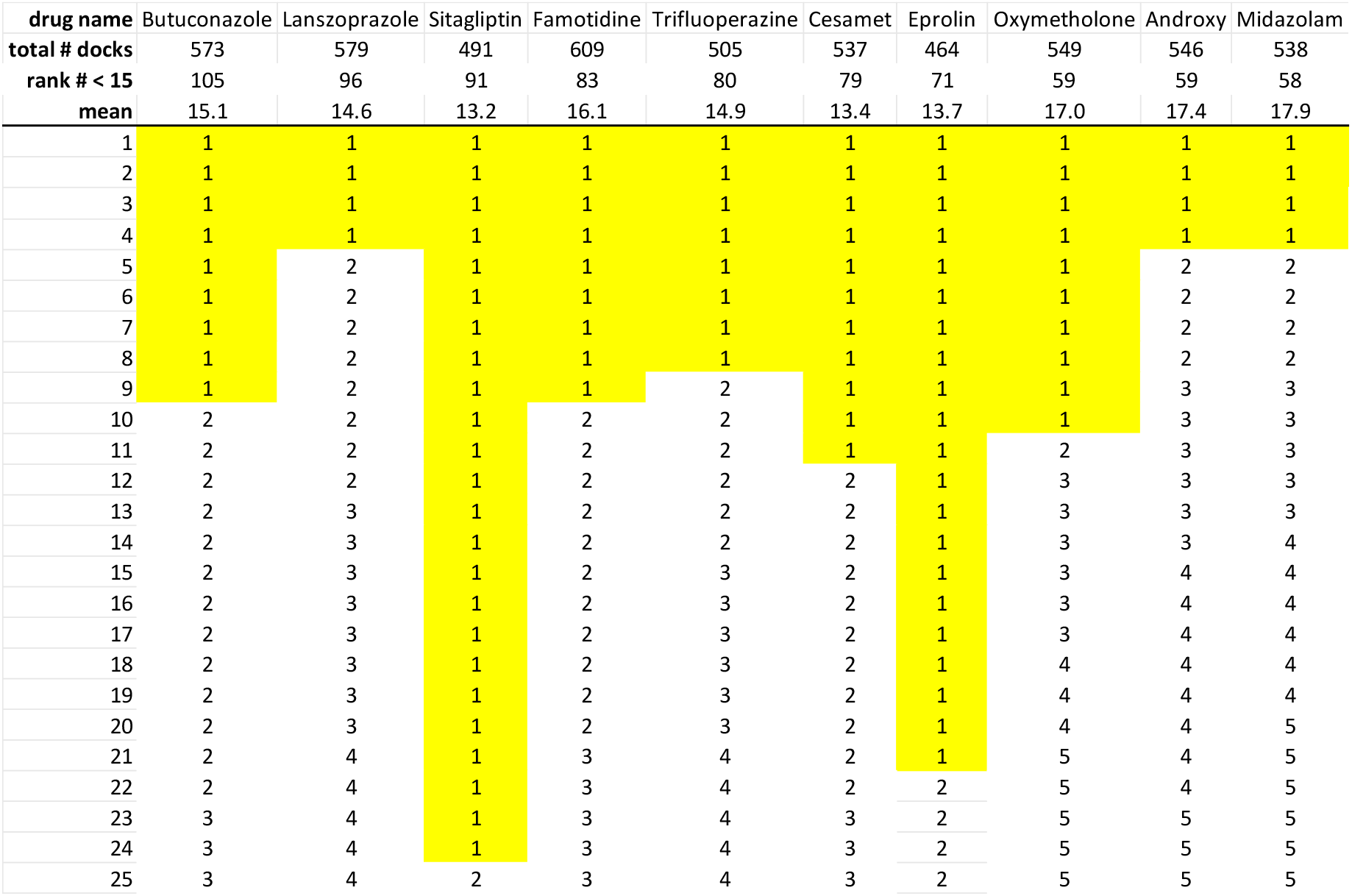
Top ten most promiscuous drugs. Yellow highlights indicate that the drug ranked #1 for a receptor.

The most promiscuous drug, butoconazole, ranked in the top 1% for 105 of the 1531 targets, while sitagliptin ranked #1 out of 24 targets (**Table 3**). The chemical structures for the top 10 drugs are illustrated in **Figure 1**. Seven out of the ten drugs display many rotatable bonds, which allow them to take on a large number of 3-dimensional poses.

On the other end of the promiscuity spectrum are the drugs with very high specificity. An ‘ideal’ drug would bind to its intended target and have low or no affinity for other receptors. **Table 4** illustrates the eight most specific drugs, and their chemical structures are shown in **Figure 2**.

**Table 4.**
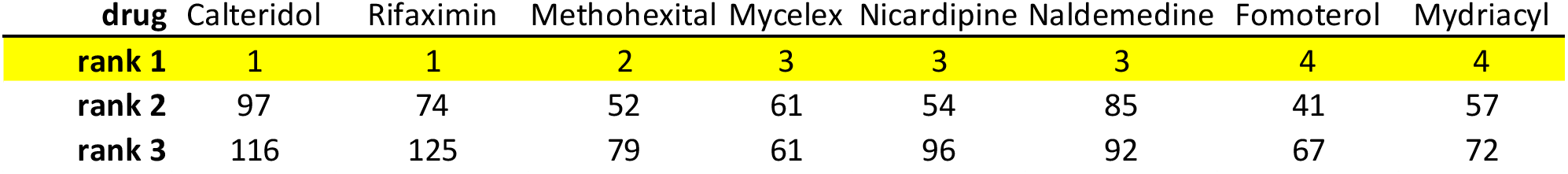
Rankings of the eight most specific drugs.

**Figure 2.**
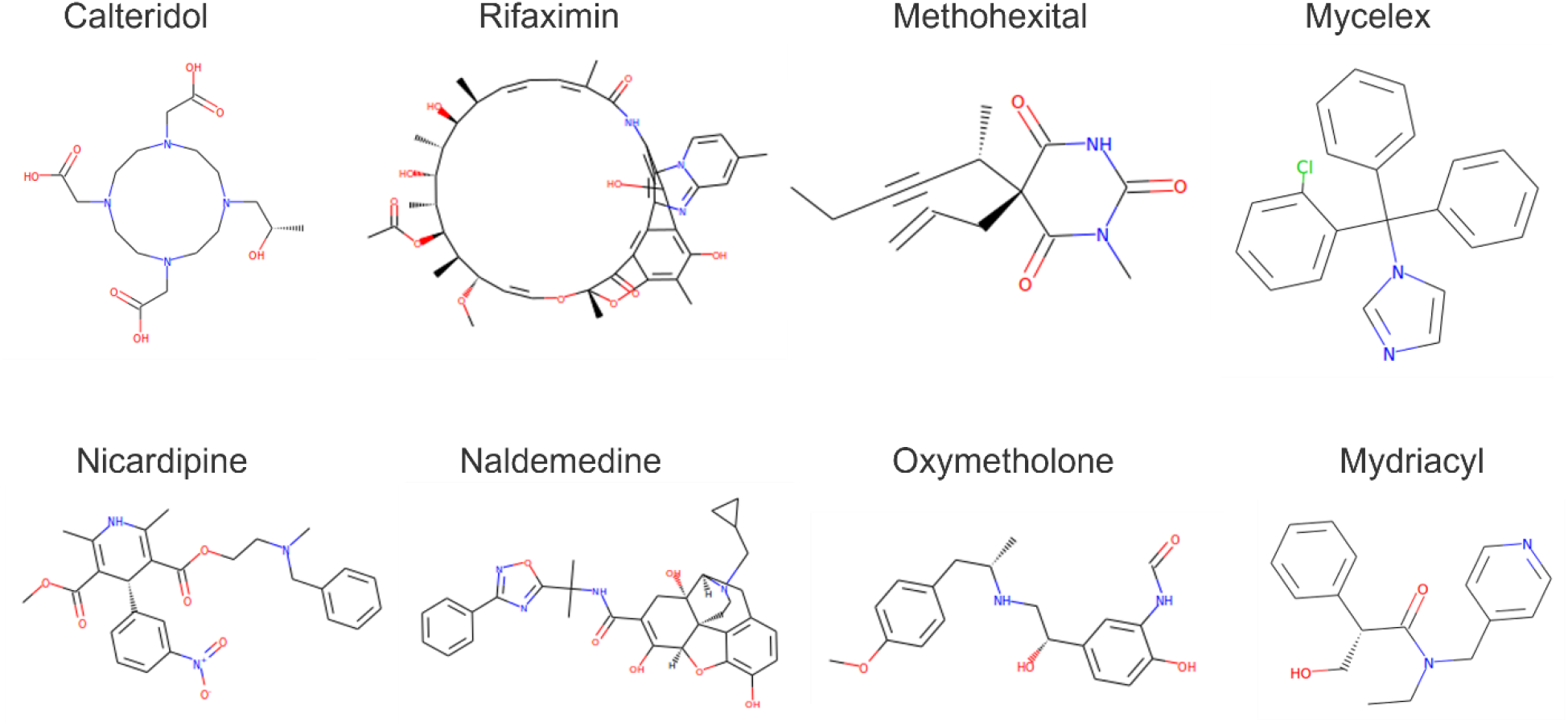
Structures of the eight most specific drugs.

The best rank for these highly specific drugs is between 1 and 4, while their second-best rank is much higher (41-97, Table 4), indicating a loss of affinity. The structures of some of these drugs help explain their high specificity: they are unusual (e.g., circular) and/or of high molecular weight.

**Figure 3** illustrates the promiscuity “rank profiles” for several drugs. It graphs the sorted rank for a drug as a function of its ranked “target number.” The profiles differ dramatically between promiscuous and specific drugs. The profile of the antifungal butoconazole is very flat as its rank remains small for a large number of receptors. The highly specific drug rifaximin, an antibiotic, rises steeply from its rank of 1 for its optimal target. The profiles for two drugs with less extreme profiles are shown as well. Lumacaftor, a cystic fibrosis drug, rises quickly initially, but then the curve becomes shallower. The rank order of nelarabine, a chemotherapy drug, is almost linearly dependent on the ranked target number. These rank profiles will be helpful in characterizing drug promiscuity.

**Figure 3.**
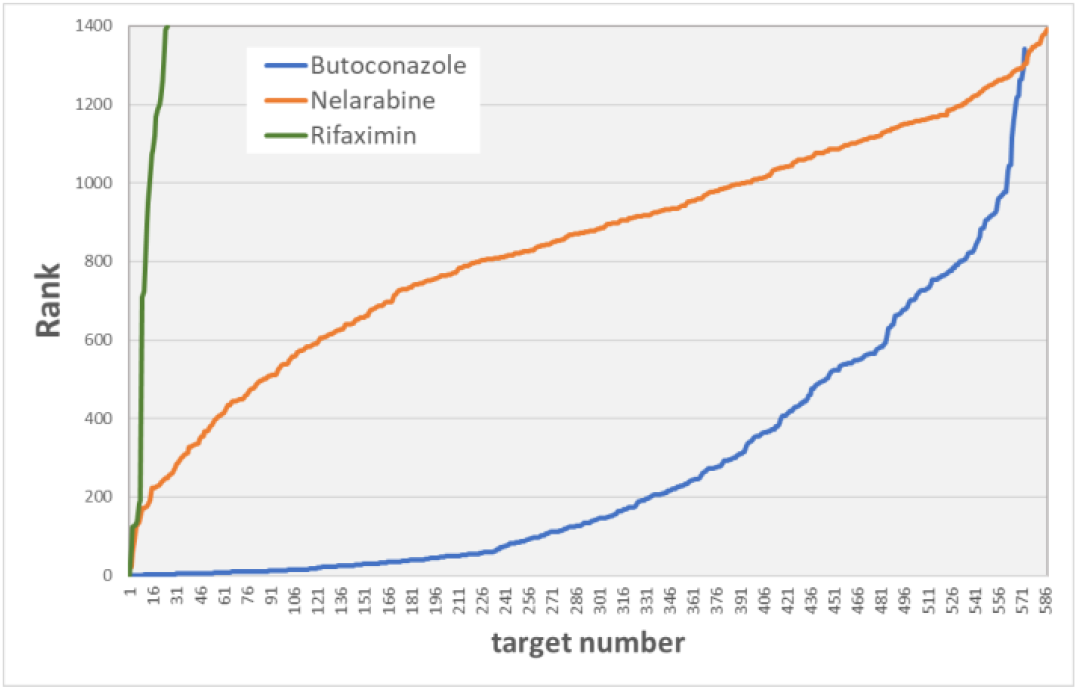
Promiscuity profiles.

Except for a few particular drugs, most small molecules bind with reasonable affinity to several target proteins. How does that affect their therapeutic performance? It depends on i) what effect drug binding has on the function of the target protein and ii) whether the target protein plays a role in the disease. If a target protein is not involved in the disease process, and the drug inhibits its function, it could cause unwanted side effects. Identifying the proteins to which a drug binds with high affinity by *in silico* docking, as shown in this paper, may shed some light on this matter. The subset of proteins analyzed here was small: only 622 were analyzed. However, the promiscuous drugs shown in **Table 3** still had enough targets to allow further study. **Table 5** illustrates the properties of the top 20 targets for the antifungal butoconazole. For each target, a literature search was performed to see whether they could play a role in fungal infection. Interestingly, 4 of the top 5 and 7 of the 20 proteins have been published as appropriate targets for antifungal therapy. These seven proteins have widely different functions, so the ability of butoconazole to bind to them with high affinity helps to explain its therapeutic efficacy in treating fungal infections.

**Table 5.**
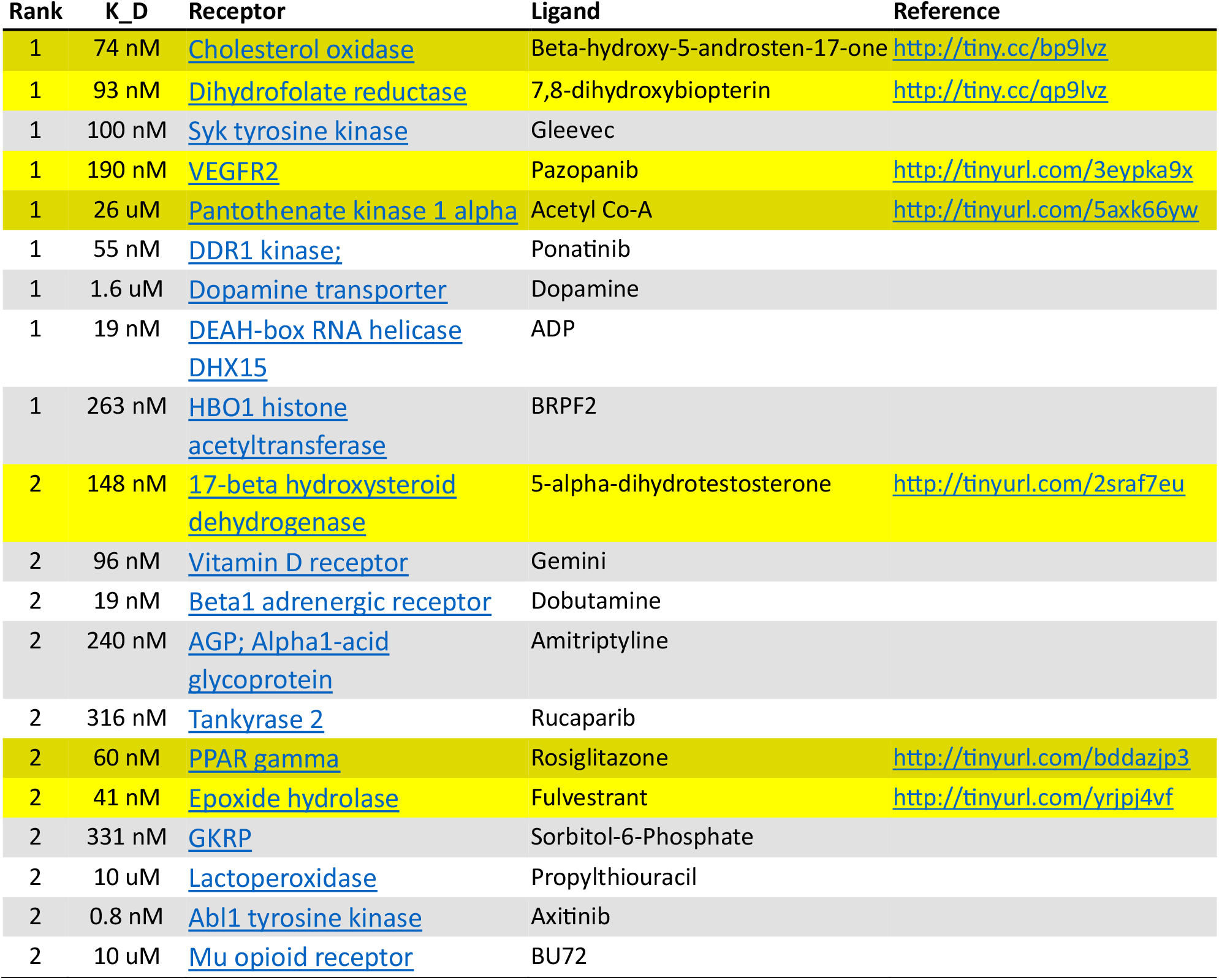
Top-ranking targets for butoconazole. List of the top 20 receptors targeted by the promiscuous antifungal butoconazole, which ranked at position 1 and 2 for 9 and 11 receptors, respectively. The Binding affinity for a receptor was estimated by the eHiTS score, which was converted to a dissociation constant KD. Both the receptor name and co-crystalized ligand are listed. Clicking on the receptor name will open the associated PDB structure page. Highlighted in yellow are seven antifungal targets. Links to the relevant papers are provided. **Acronyms**: VEGFR=Vascular Endothelial Growth Factor, DDR1=Discoidin Domain Receptor 1, PPAR=Peroxisome Proliferator Activated Receptor, GKRP=GlucoKinase Regulatory Protein.

## Discussion

In this paper, I analyzed the interaction between 1531 FDA-approved drugs and 622 proteins co-crystalized with a drug or neurotransmitter, using *in silico* docking of the drugs into the binding site of the co-crystalized ligand. For the 952,282 possible combinations, 624,498 were successfully docked, indicating that 327,784 drugs were incompatible with the targeted binding site. Either because they were too large or because of a steric hindrance problem. For each drug, the binding affinity of the optimal poses was estimated by the eHiTS software and used to rank the target proteins (receptors). Each drug now has a ranking for each compatible receptor. Note that this ranking is relative to all other drugs binding to the same protein. The relative rankings for a drug were sorted, and the top 1% of the receptors with rankings below 15 were identified. **Table 3** lists twenty highly promiscuous drugs that ranked better than 1% for 58-105 receptors.

This analysis also allowed the identification of highly specific drugs that bound to only one receptor with high affinity (**Table 4**). There were 278 drugs (18%) with this extreme specificity. Their chemical structures (**Fig. 2**) provide some clues that can help explain the unusual specificity. A sizeable molecular weight may exclude them from all but the largest binding sites. Ring shapes and other steric confinements may limit binding to unique compatible pockets. Most drugs do not belong to either of these two extreme classes. However, 82% of the drugs are binding with high affinity to at least two receptors. But only 622 receptors were analyzed here. If we extrapolate this finding to the ∼25,000 proteins expressed in the human body, most drugs will target at least 100 receptors. Therefore, drug promiscuity is a universal problem. Promiscuity profiles (**Fig. 3**) may be helpful in designing a suitable classification system. This idea needs to be further developed.(Nakane 2020)

The most significant challenge going forward will be to interpret the promiscuity data. Which of the identified receptor targets enhance the therapeutic efficacy of a drug, and which targets are responsible for unwanted side effects. Consider the targets for one of the most promiscuous drugs, butoconazole, an antifungal compound (**Table 5**). Four of the five top receptor proteins are antifungal targets. These will likely contribute to the therapeutic efficacy. Their functions are distinct: an oxidase, a reductase, two kinases, and a receptor. Identifying such diverse therapeutic targets may help identify the mechanism of action for a drug. The fact that the top twenty receptors for butoconazole contained seven antifungal targets validates the docking procedure.

Another challenge that lies ahead is increasing the number of proteins to more closely resemble the repertoire expressed in the human body. Not all proteins have been crystallized. However, single-particle cryo-EM methods have reached atomic resolution (Nakane 2020). They rapidly increase the number of available protein structures, including many membrane proteins inaccessible to X-ray crystallization. Another recent development is AlphaFold, an AI system developed by DeepMind that predicts a protein’s 3D structure from its amino acid sequence (https://alphafold.ebi.ac.uk/). These structures may turn out to be useful for drug discovery.

The majority of protein structures are not co-crystallized with a ligand, so a suitable binding pocket needs to be identified. Some docking software packages can identify binding pockets in protein structures (ICM-Pro, MolSoft LLC, San Diego), but the accuracy of this approach remains to be seen. It also increases the computational workload. The challenges identified above indicate that scaling up the approach used here to characterize drug promiscuity will require a significant investment in time and resources. However, the results obtained in this small-scale study suggest it will be worthwhile.

## Notes

### Competing Interest Statement

The authors have declared no competing interest.

## References

Alper, A. B., Jr., H. Tomlin, U. Sadhwani, A. Whelton and J. Puschett (2006). “Effects of the selective cyclooxygenase-2 inhibitor analgesic celecoxib on renal carbonic anhydrase enzyme activity: a randomized, controlled trial.” Am J Ther 13(3): 229–235.

Irwin, J. J., T. Sterling, M. M. Mysinger, E. S. Bolstad and R. G. Coleman (2012). “ZINC: a free tool to discover chemistry for biology.” J Chem Inf Model 52(7): 1757–1768.

Irwin, J. J., K. G. Tang, J. Young, C. Dandarchuluun, B. R. Wong, M. Khurelbaatar, Y. S. Moroz, J. Mayfield and R. A. Sayle (2020). “ZINC20-A Free Ultralarge-Scale Chemical Database for Ligand Discovery.” J Chem Inf Model 60(12): 6065–6073.

Nakane, T., A. Kotecha, A. Sente, G. McMullan, S. Masiulis, P. Brown, I. T. Grigoras, L. Malinauskaite, T. Malinauskas, J. Miehling, T. Uchanski, L. Yu, D. Karia, E. V. Pechnikova, E. de Jong, J. Keizer, M. Bischoff, J. McCormack, P. Tiemeijer, S. W. Hardwick, D. Y. Chirgadze, G. Murshudov, A. R. Aricescu and S. H. W. Scheres (2020). “Single-particle cryo-EM at atomic resolution.” Nature 587(7832): 152–156.

Zsoldos, Z., D. Reid, A. Simon, B. S. Sadjad and A. P. Johnson (2006). “eHiTS: an innovative approach to the docking and scoring function problems.” Curr Protein Pept Sci 7(5): 421–435.

Zsoldos, Z., D. Reid, A. Simon, S. B. Sadjad and A. P. Johnson (2007). “eHiTS: a new fast, exhaustive flexible ligand docking system.” J Mol Graph Model 26(1): 198–212.

